# Direct comparison of constitutive Rax-Cre transgenic drivers that activate in the mouse embryonic eye field

**DOI:** 10.1101/2025.10.11.681822

**Authors:** Nadean L. Brown, Samuel Goodyear-Brown, Sabine Fuhrmann

**Affiliations:** Department of Cell Biology & Human Anatomy, University of California, Davis, CA; Department of Ophthalmology and Visual Sciences, Vanderbilt Eye Institute, Vanderbilt University Medical Center, Nashville, TN; Department of Cell and Developmental Biology, Vanderbilt University Medical School, Nashville, TN

**Author notes:** The authors declare no competing interests and nothing to disclose.

**Keywords:** Rax, retinal development, RPE, Cre recombinase, transgenic mice

## Abstract

**Purpose:** Multiple mouse lines using constitutive or inducible Cre recombinase expression have taken advantage of the early optic field expression of the Rax/Rx gene and its subsequent, progressive restriction to retinal progenitors, retinal pigmented epithelium and the optic stalk. Among these, the constitutive transgenic lines, Rx3-Cre and Rax-Cre BAC Tg, are currently used by vision researchers. Here we directly compare their prenatal ocular and extra-ocular Cre activities.

**Methods:** Rx3-Cre or Rax-Cre BAC transgenic male mice were mated to the Cre-dependent lineage tracer Ai9/tdTomato. The resulting live and fixed tissue expression for each line was evaluated at multiple stages of development, from the optic vesicle stage onward. Both whole embryo and immunolabeled cryosectioned material were digitally imaged.

**Results:** Rx3-Cre recapitulates endogenous RAX protein expression at the optic vesicle and optic cup stages, but there is also ectopic Cre activity in periocular mesenchyme (POM), within PITX2- and PECAM-1-expressing subpopulations. We also show that ectopic Cre activity can include germline expression. Besides a small ectopic domain in developing limbs, the Rax-Cre BAC mouse driver fully recapitulates endogenous RAX expression in the embryonic and postnatal eye.

**Conclusions:** We provide a systematic analysis of two Rax-Cre drivers during embryonic and postnatal eye development that are very useful to recapitulate severe congenital eye diseases. Data presented here strongly support the inclusion of lineage reporting and evaluation of littermates lacking a Cre transgene, whenever conditional targeting strategies are used in studies of optic vesicle-derived tissues.

## Introduction

Eye formation is a multi-step, dynamic process that is tightly regulated by a complex gene regulatory network^1, 2^. In vertebrates, a central eye field is specified in the anterior neural plate that splits into bilateral optic vesicles. Multiple factors then regionalize the growing optic vesicles, and subdivide them into optic stalk (OS), neural retina (NR) and retinal pigment epithelium (RPE) tissues. Both ocular and extraocular tissues are integral during tissue regionalization and eye morphogenesis. For example, the NR interaction with adjacent embryonic surface ectoderm triggers optic vesicle invagination into an optic cup. The outer optic cup cells also receive input from surrounding peri-ocular mesenchyme (POM) to differentiate into RPE, while inner optic cup cells receive NR-promoting signals from the surface ectoderm to commit to neuroepithelial fate. The gene networks driving these processes are critical since individual genetic mutations result in partial or missing eyes. Thus, better mechanistic understanding of the molecular steps of eye development is necessary for advancing clinical treatments that includes modeling it using *in vitro* stem cell and/or organoid technologies. One gene whose expression is activated in the eye field, prior to bilateral eye formation, is the homeobox transcription factor *Rax/Rx*^3, 4^. During human first trimester development, *RAX* is essential for eye formation, and its loss induces anophthalmia or microphthalmia^3–8^. Further evidence that the *Rax* gene rests at the top of an eye development gene hierarchy, is the finding that its loss in frogs, fish, or axolotls also causes eyelessness^6, 9^.

The discovery and application of site-specific recombinases revolutionized mouse models as tools for investigating ocular development and disease. By driving the expression of microorganism-derived Cre recombinase in a spatially-specific manner, the early lethality of many developmental genes was circumvented. The advent of inducible Cre drivers further expanded the conditional mutant toolkit by adding the control of timing and duration for the induction of tissue-specific mutations. The first mouse Rax/Rx Cre line, Tg(rx3-icre)1Mjam (hereafter Rx3-Cre), used an evolutionary conserved 4kb segment of noncoding DNA, located 5’ upstream of the *Oryzias latipes* (medaka) Rx3 gene promoter to drive Cre recombinase^10^. Rx3-Cre has robust activity in the eye field, NR, RPE and OS; and its conditional removal of a *Rax^flox/flox^* allele phenocopied *Rax^-/-^*germline mutants^10^. For these reasons Rx3-Cre mice were used to recombine at least 37 conditional alleles (MGI database: www.informatics.jax.org).

However, there are increasing reports of ectopic expression in the lens, neural crest and/or olfactory epithelium, and most concerning, ectopic germline activity. Thus, mutant studies using Rx3-Cre mice could produce inconsistent and potentially confounding outcomes, particularly for extracellular signaling genes that act broadly and/or reiteratively during eye development.

Another constitutively active Rax-Cre line is a mouse BAC transgenic, made by the GENSAT project and cryobanked at the MMRRC^11^. This strategy was also used to create an independent Rax-Cre BAC transgenic mouse line^12^. We cryo-revived the GENSAT transgenic line, tested the fidelity of its expression and subsequently used it to generate conditional mutant phenotypes for several genes^13, 14^. Rax-Cre BAC Tg mice display uniform activity throughout the eye field and its derivatives. Here we provide additional, relevant information about the Rax-Cre BAC transgenic lineage, directly comparing it to the Rx3-Cre driver, during the onset and progression of eye development.

## Materials and Methods

### Animals

Mouse strains used in this study are Gt(ROSA)26Sor^tm9(CAG-tdTomato)Hze^ (Ai9) Cre lineage tracer maintained on a C57BL/6 background (JAX#007909)^14^;^11^; Tg(CAG-Bgeo/GFP)21Lbe/J (Z/EG) (JAX#003920); Rx3-Cre transgenic insertion (Tg(rx3-icre)1Mjam), Nf2^tm2Gth^; Wwtr1^tm1.2Hmc^Yap1^tm1.2Hmc^/WranJ (Jax#030532), B6.C-Tg(CMV-cre)1Cgn/J (Jax#006054); Rax-Cre BAC transgenic line (Tg(Rax-Cre) NL44Gsat/Mmucd cryobanked at MMRRC UC Davis (Stock Number: 034748-UCD). Rax-Cre BAC mice were cryopreserved on a mixed coat color background but subsequently maintained by backcrossing to CD-1. For all experiments, genotyping was performed as described^11, 13, 15^ or by Transnetyx (Cordova, TN) using Taqman and custom-designed probes. Noon of the vaginal plug date was assigned to age E0.5. All mice were housed and cared for in accordance with guidelines provided by the National Institutes of Health and the Association for Research in Vision and Ophthalmology, with oversight from the UC Davis and VUMC Institutional Animal Care and Use Committees.

### Immunofluorescent Labeling

Embryonic heads were fixed in 4% paraformaldehyde/PBS for 1 hour on ice or at RT, processed by stepwise sucrose/PBS incubations, and embedded in Tissue-Tek OCT. Cryostat sections (10-12 μm) were labeled with Goat anti-tdTomato/DsRed (1:500 or 1:1,000, SicGen), Chick anti-GFP (1:2000, Aves), Rabbit anti-Rax (1:1000, Takara), Rabbit anti-PECAM-1 (1:150, Becton Dickinson) or Rabbit anti-Pitx2 (1:1,000, Capra Science), followed by incubation with secondary antibodies: Donkey anti-Goat Alexa594 (1:1000, Jackson Immunologicals), Donkey anti-Goat Rhodamine Red (Jackson ImmunoResearch), anti-Chick IgY Alexa488 (1:2000, Invitrogen), Donkey anti-Rabbit Alexa488 (1:500, Jackson Immunologicals), Donkey Anti-Rabbit Alexa Fluor647 (1:1,000, Jackson ImmunoResearch).

Nuclei were counterstained with DAPI and slides mounted with Fluorosave (MilliporeSigma) or Prolong Gold Antifade (Invitrogen). For each age and marker, 4 embryos with either Cre transgene and the Ai9 reporter were analyzed. To track germline transmission, we generated Rx3-Cre;Ai9 embryos with *Nf2*, *Yap* and *Taz* alleles.

### In situ hybridization

Whole mount in situ hybridization was performed on wild type CD1 embryos or isolated limbs as described in^16, 17^ using a 5’ mouse Rax/Rx cDNA antisense digoxygenin labeled riboprobe^4^. Embryos, but not limbs, were digested with proteinase K prior to hybridization.

### Microscopy

Some living embryos with td-Tomato expression were imaged using a Leica MZ12 dissecting microscope, equipped with a UV light source, Spot camera and software (v5.2) or a Leica DFC7000T camera and LAXS software (v3.7.5). Other embryos were imaged after fixation with an AxioZoomV16 with Hamatsu Fusion camera (HDCamC14440-20UP), Nikon SMZ1270 epifluorescence stereomicroscope equipped with DS-Fi3 camera and Nikon NIS Elements BR 5.11.03 (Build 1373) software, and an Olympus SZX12 stereomicroscope with Microfire camera and PictureFrame software. Antibody-labeled cryosections were imaged using a Leica DM5500 microscope, equipped with a SPEII solid state laser scanning confocal optic and processed using Leica LASX (v.5), or a ZEISS LSM880 and processed using Zen v2.3 SP1 (Black), FIJI/Image J (NIH) and Adobe Photoshop (CS5, 26.10.0 versions) software programs.

## Results

To date, at least eight Rax transgenic mouse lines have been created, characterized and described in publications (Table 1). The first, a Rax BAC Tg GFP reporter has been an important tool in the development of mouse ES and organoid experimental strategies, to mark and follow the formation of the optic vesicle/cup in culture paradigms^12^. Table 1 summarizes published information about Cre drivers that used Rax/Rx gene sequences to drive Cre expression/activity, both available or extinct. In this paper we compare the activities of two constitutive Rax mouse lines, highlighting the fidelity of a recombineered Rax BAC transgene.

**Table 1.**
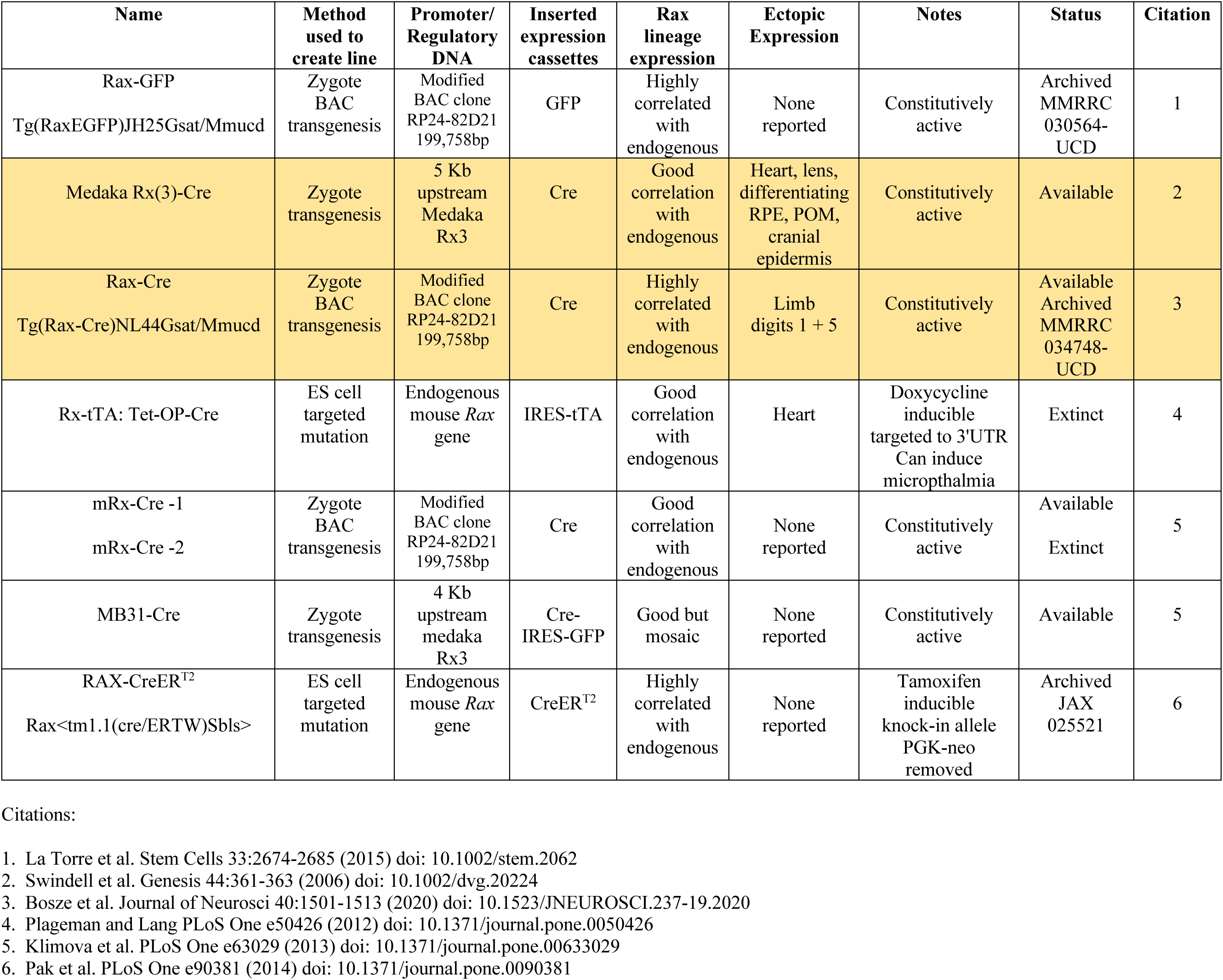
Comparison of Rax/Rx-Cre mouse models.

### The Rx3-Cre mouse driver

We used Rx3-Cre to manipulate *Porcn* and *Nf2* gene expression during optic cup morphogenesis and observed ectopic reporter expression in lens ectoderm and periocular mesenchyme (POM)^18, 19^. Moreover, in a separate, ongoing study a more systematic analysis revealed that Rx3-Cre can induce germline deletion (see below). However, here we systematically re-evaluated Rx3-Cre activity using a Cre-inducible Ai9 (tdTomato) expression, versus endogenous RAX protein detection, and the identification of POM cell types with ectopic Rx3-Cre activity.

Previous studies reported that Rx3-Cre activity starts in the eye field at E8.75-E9.0^10, 12^ (SF, unpublished observations) and continues in the early optic vesicle with expression domains in NR, RPE, OS, lens vesicle and subset of POM cells^12, 18, 19^. Here we interrogated Rx3-Cre activity in embryos that are also heterozygous for Ai9, or in more complex genotypes that combined Rx3-Cre and Ai9 with *Nf2*, *Yap* and/or *Taz* heterozygotes. We confirmed Rx3-Cre-induced tdTomato expression in the optic vesicle, ventral forebrain and heart (Fig. 1A). Anti-DsRed detection of the TdTomato protein revealed additional Cre activity in epidermal cells covering the head including the surface ectoderm, and in a subpopulation of POM cells (Fig. 1A’). The POM originates from neural crest and mesodermal lineages that differentiate into distinct extraocular cell populations giving rise to vascular structures, choroid and sclera, hyaloid vasculature, and components of the anterior eye such as the corneal stroma and endothelium, iridocorneal angle, ciliary muscle and iris^20, 21^. At E10.5, tdTomato is expressed within the optic cup (presumptive retina and RPE), lens vesicle, adjacent diencephalic neuroepithelium, ventral telencephalic neuroepithelium, cranial mesenchyme including POM, and in heart (Fig. 1B, B’).

**Figure 1.**
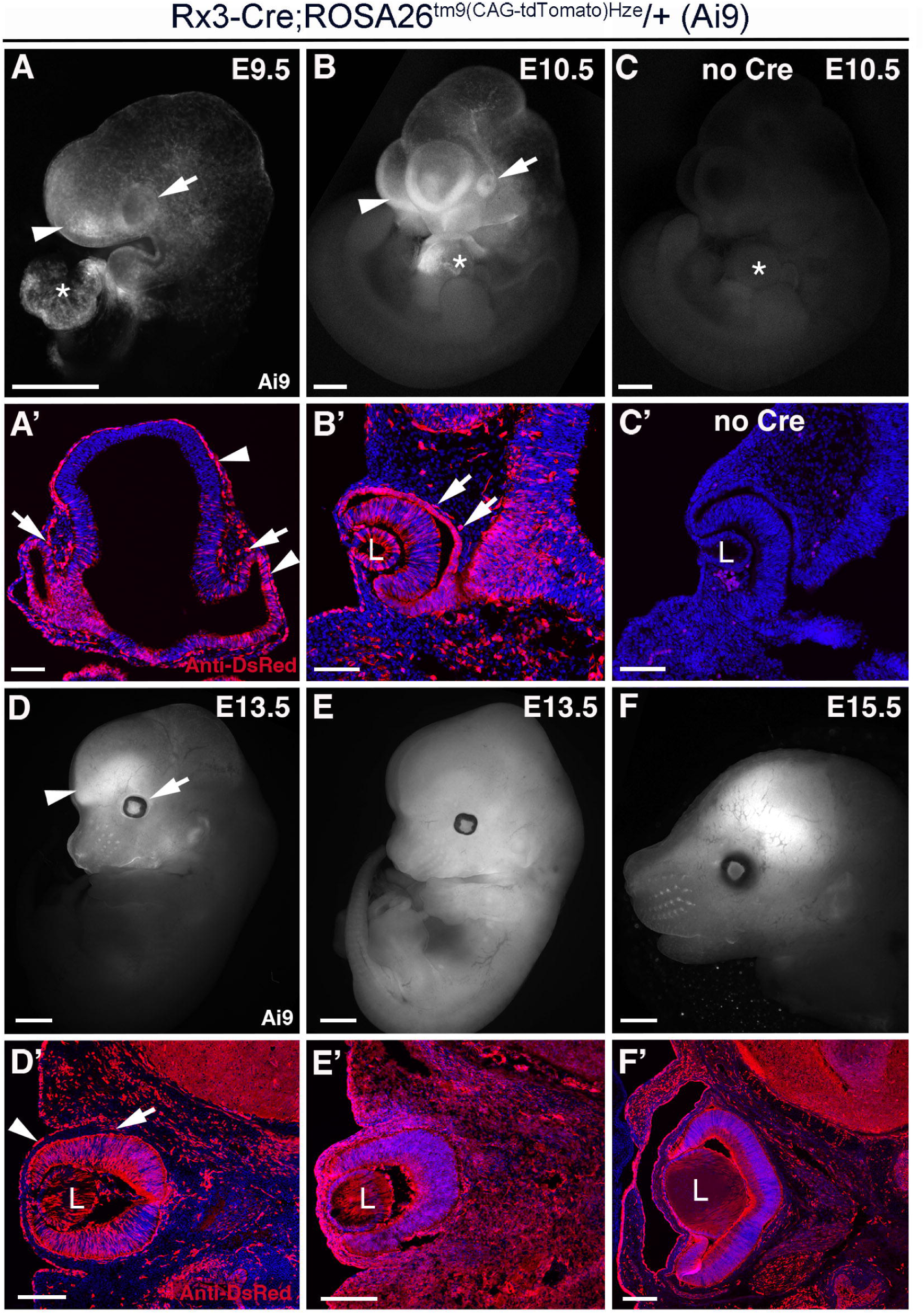
Rx3-Cre activity in optic vesicle-derived tissues, plus surface ectoderm and periocular mesenchyme between E9.5 and E15. A-F) Ai9 (tdTomato) reporter expression in the optic vesicle/cup (arrow), surrounding cranial regions including telencephalon (arrowhead), adjacent diencephalon and periocular mesenchyme, and in the heart (asterisk) at E9.5 (A) and E10.5 (B). No reporter expression is observed in the absence of Rx3-Cre (C, C’). At E13.5, tdTomato expression is widespread in the head, particularly in ocular (arrow) and extraocular tissues (arrowhead, D). In (E), an example for globally activated Rx3-Cre is shown. At E15.5, Rx3-Cre activity also labeled cranial tissues including the eye (F). **A’-F’)** Coronal sections labeled with anti-DsRed antibody through forming eyes of embryos in A-F highlight ocular and extraocular Rx3-Cre activity in head and surface ectoderm (arrowheads) and POM (arrows). L = lens, * = heart. Scale bars A, B, C = 0.5mm; D, E, F = 1mm; A’ - C’= 100 microns; D’ - F’ = 200 microns.

There is no tdTomato expression without Rx3-Cre (Fig1 C, C’). At E13.5, the Rx3-Cre transgene induced tdTomato expression in the retina, RPE and lens, and adjacent diencephalic neuroepithelium, cranial epidermis and in a subset of POM and adjacent cranial mesenchyme cells (Fig. 1D, D’). Occasionally, we observed embryos with Ai9 expression throughout the body, indicating global Rx3-Cre activity (Fig. 1E, E’). Specifically, among the 30 Rx3-Cre-positive embryos analyzed in this study, we observed 2 embryos with globally-activated tdTomato expression (6.7%). At E15.5, tdTomato expression is essentially the same as that described for E13.5 (Fig. 1F, F’).

Globally activated Ai9 reporter expression may include the germ line. Indeed, analysis for excised alleles of 3 exemplary genes (*Taz, Yap, Nf2*; separate ongoing project) confirmed germline deletion of *Taz* in 29.3% of embryos without Rx3-Cre (n=99), *Yap* in 5% (n=99) and *Nf2* in 12.1% of embryos without Rx3-Cre (n=99). We also assembled a pedigree showing transmission of fully excised (germline) *Taz* alleles in Rx3-Cre-negative offspring in generation IV (Suppl Fig. 1).

To compare Rx3-Cre-induced tdTomato reporter expression with endogenous Rax protein, we co-labeled cryosections with DsRed and RAX antibodies (Fig. 2). At E9.5 and E10.5, RAX protein is detectable in the distal and proximal optic vesicle and in neuroepithelial cells of the adjacent diencephalon (Fig. 2A,B). In contrast, Rx3-Cre-mediated recombination induced ectopic tdTomato in the developing lens, differentiating RPE and in a subset of the POM (Fig.2A’-D’). The merged labeling shows the extent of ectopic Rx3-Cre activity (Fig.2A“-D”), particularly in the POM. The homeodomain transcription factor *Pitx2* is required for eye development and expressed in the neural crest-derived lineage of the POM^20^. Analysis of PITX2 expression showed that a subset of Rx3-Cre-recombined TdTomato-positive cells is robustly co-labeled for PITX2 at E10.5 and E13.5 (Fig. 3A-D’). A proportion does not express PITX2, thus, we co-labeled with the endothelial marker PECAM-1 that is primarily present in mesodermally-derived cells^22^. Interestingly, we observed POM cells directly lining the basal side of the optic cup (RPE) co-labeled for both TdTomato and PECAM-1 at E10.5 and E13.5 (Fig. 3E-H’). In summary, these observations suggest that Rx3-Cre exhibits recombination activity in several extraocular domains of different origins starting in the early optic vesicle.

**Figure 2.**
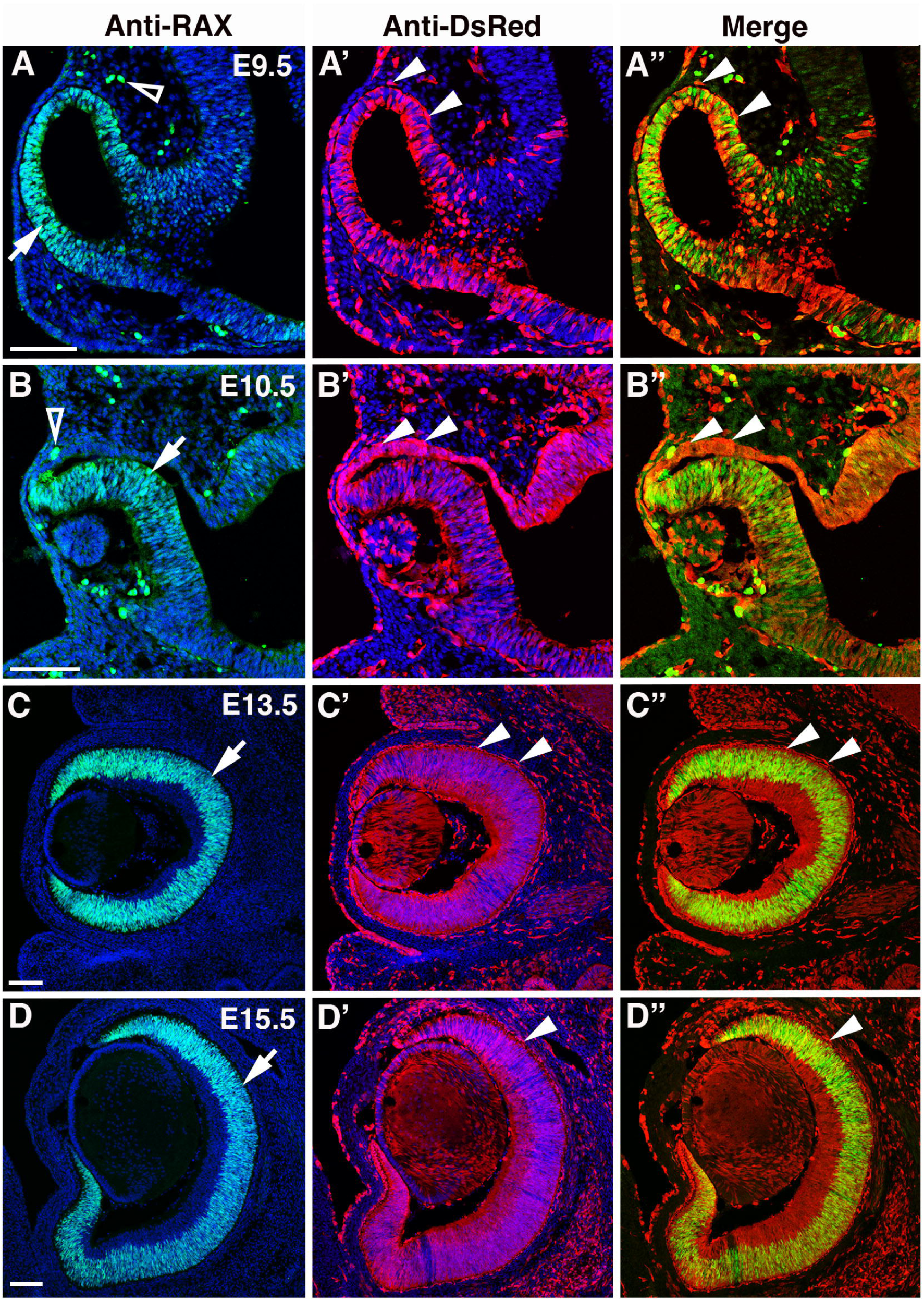
Direct comparison of endogenous Rax protein and Rx3-Cre;tdTomato expression. A-A”) Endogenous Rax is present in the E9.5 ocular neuroepithelium (A, arrow), whereas tdTomato+ expression is also evident in adjacent brain neuroepithelium, surface ectoderm and POM cells (A’, A”, arrowheads). **B-D”).** Open arrowheads mark vascular cells with autofluorescence (A, B). At E10.5 - E15.5, Rax is expressed by retinal progenitor cells (B-D, arrows), but tdTomato is also present in the lens, epidermis and POM cells (C’-D”, arrowheads). **D-D”).** Rax is localized in retinal progenitor nuclei, but the tdTomato-only lineage includes differentiated RGCs in forming GCL. OV = optic vesicle, L = lens. Scale bars = 100 microns.

**Figure 3.**
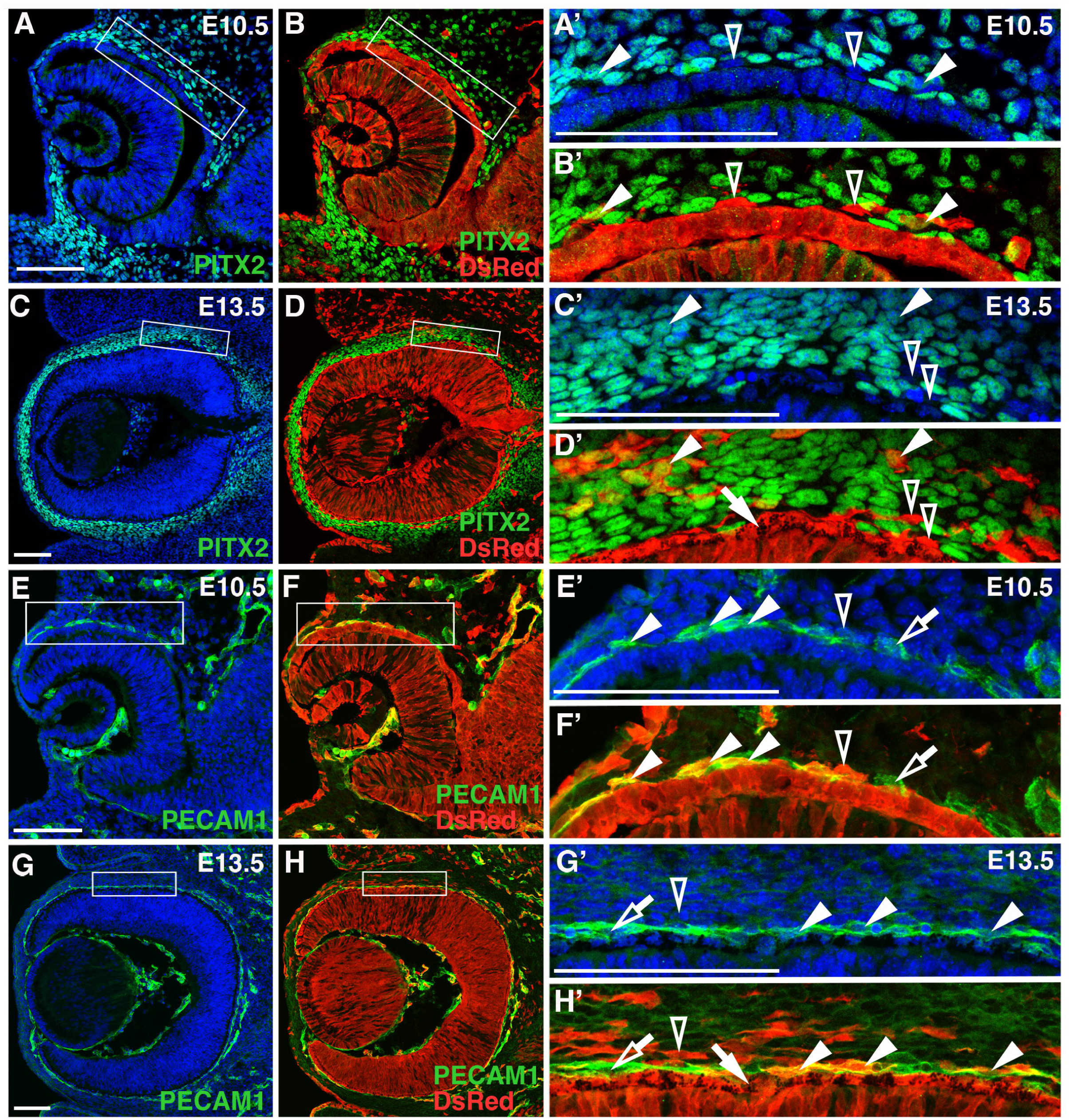
Rx3-Cre-tdTomato activity in the periocular mesenchyme is detectable in neural crest-derived and endothelial lineages. A-D) At E10.5 and E13.5, the Rx-3 Cre lineage (anti-DsRed) includes the neural crest-derived POM (PITX2+, green). Boxed areas in A-C are shown magnified in A’ - D’. Several examples of tdTomato+PITX2+ mesenchymal cells (filled arrowheads), and tdTomato-only cells (open arrowheads) (A’-D’). **E-H)** PECAM-1 expression (green) marks endothelial cells in the POM and vitreal space. Here too are TdTomato+PECAM-1+ (filled arrowheads; E’-H’), but also tdTomato-only cells (open arrowhead), and PECAM-1 only POM cells (open arrows). In D’ and H’, the pigmented RPE layer (filled arrow) is labeled with Anti-DsRed/tdTomato. Scale bars = 100 microns.

### The Rax-Cre BAC mouse driver

Three different lines have been generated using the mouse BAC clone RP24-82D21 containing 199,758 nucleotides of Chromosome 18 with the mouse *Rax* gene centrally located within it. The GENSAT/MMRRC project used bacterial recombineering to insert Cre or GFP expression cassettes, and pronuclear injection of the recombineered DNA, to establish stable transgenic lines^11^. An independent effort with this BAC Clone also generated the mRx-Cre BAC transgenic line^12^. We resuscitated the Tg(Rax-Cre)NL44Gsat line from the UC Davis MMRRC repository and maintain it by backcrossing to CD1 mice, which generate robust litters and facilitate live imaging of the eye (Figs 4,5). When Rax-Cre Tg mice are mated to those carrying either Ai9 (tdTomato) or Z/EG (GFP), the fluorescent reporters show which cells and tissues have Cre activity. We observed Rax-Cre initiates recombination during gastrulation at E8, marking the eye fields in the optic sulcus (Fig. 4A) and the optic vesicles by E9 (Figs 4B,4C).

**Figure 4.**
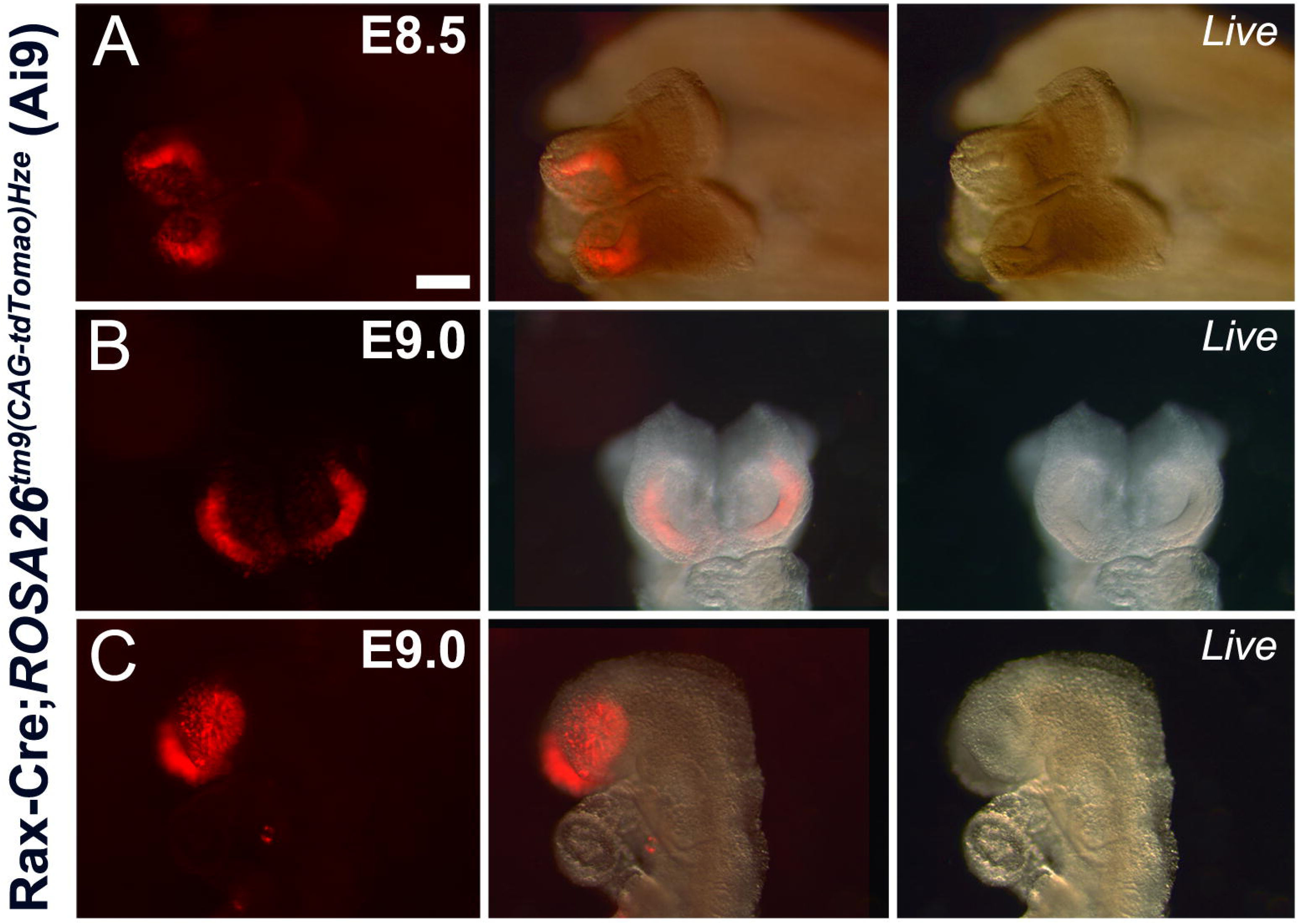
Rax-Cre BAC Tg expression in the developing eye field. A-C) Cre activated Ai9 (tdT-Tomato) live fluorescence in the optic sulcus of E8 embryos (A), and optic vesicles at E9.0 (B,C). Left panel is live fluorescence, middle is the merged image and right is brightfield illumination. Scale bar A-C = 3mm

**Figure 5.**
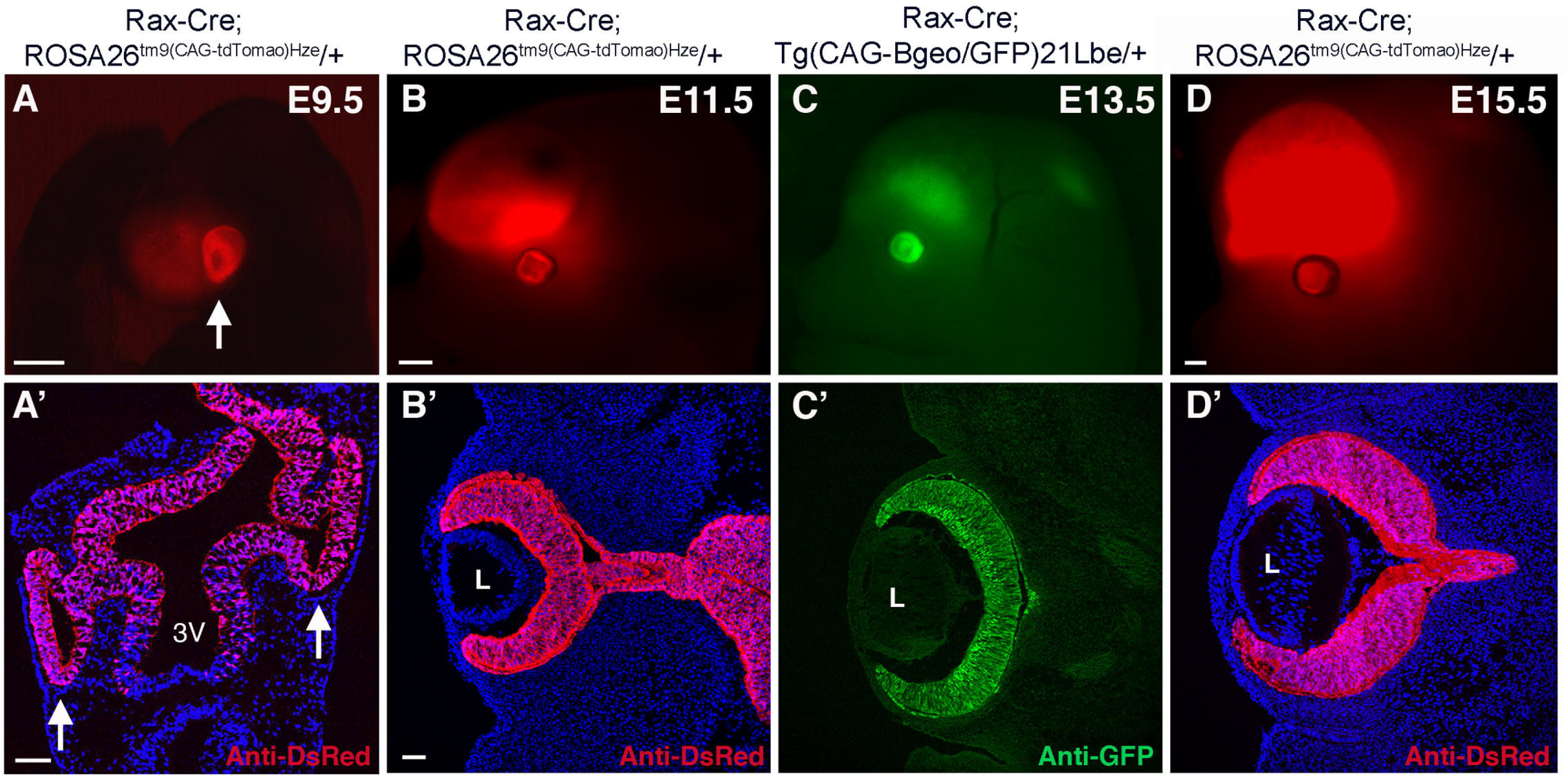
Rax-Cre BAC Tg lineage includes optic vesicle derived tissues. A-D) Live images of mouse embryos showing tdTomato expressing ocular cells (arrows in A,A’) and underlying brain domains. **A’-D’)** Horizontal sections labeled with anti-DsRed or anti-GFP through forming eyes of embryos in A-D highlight specificity of Cre activity to the optic vesicle derived tissues. L = lens, 3rd = third ventricle. Scale bars A = 1mm; B,C = 2mm; D = 1mm; A’ = 75 microns; B’-D’ = 50 microns.

Beginning at E9.5, we noted ocular and developing telencephalon tdTomato+ expression (Figs 5A,A’). Both domains persist throughout perinatal development (Figs 5,8), with the forebrain domain expanding as the cortex grows. Endogenous *Rax* mRNA and protein are also present in these same domains ^23–25^. E9-E15 cryosections show no reporter mosaicism in the cortex, ventral thalamus, optic vesicle, and its derivative tissues, the retina, RPE, and optic stalk. (Figs 5,6, data not shown). Next we colabeled E9 - E13 cryosections from Rax-Cre Tg/+; Ai9/+ embryos with specific antibodies for tdTomato/DsRed and endogenous Rax protein (Fig. 6). We note highly correlated coexpression in the optic vesicle (OV), but more prominent TdTomato expression in adjacent brain domains (Figs 5A’,6A-6A”). Starting at E11, endogenous Rax protein expression is restricted to optic cup/ retinal progenitors, highlighting tdTomato-only expression in the adjacent RPE (Fig. 6B-6B”). At this age, we also observe many more tdTomato+ cells in the hypothalamic progenitors surrounding the 3rd brain ventricle (Fig. 6C-6C”), when endogenous Rax expression is confined to the ventral brain and forming Rathke’s’ pouch (arrow in C). This precursor of the posterior pituitary also endogenously expresses *Rax*. The *Rax* gene is also normally found expressed in the developing hindbrain, with progressive temporal restriction within the hippocampus. Here too, we found tdTomato expression to be highly correlative (not shown). At E13.5 endogenous Rax protein is specific for retinal progenitor cells (Fig. 6D-6D”), but tdTomato expression remained uniform, within the RPE, optic stalk (not shown) and retina, including differentiating retinal neurons (Fig. 5D’, red only cells in Fig. 6D”).

**Figure 6.**
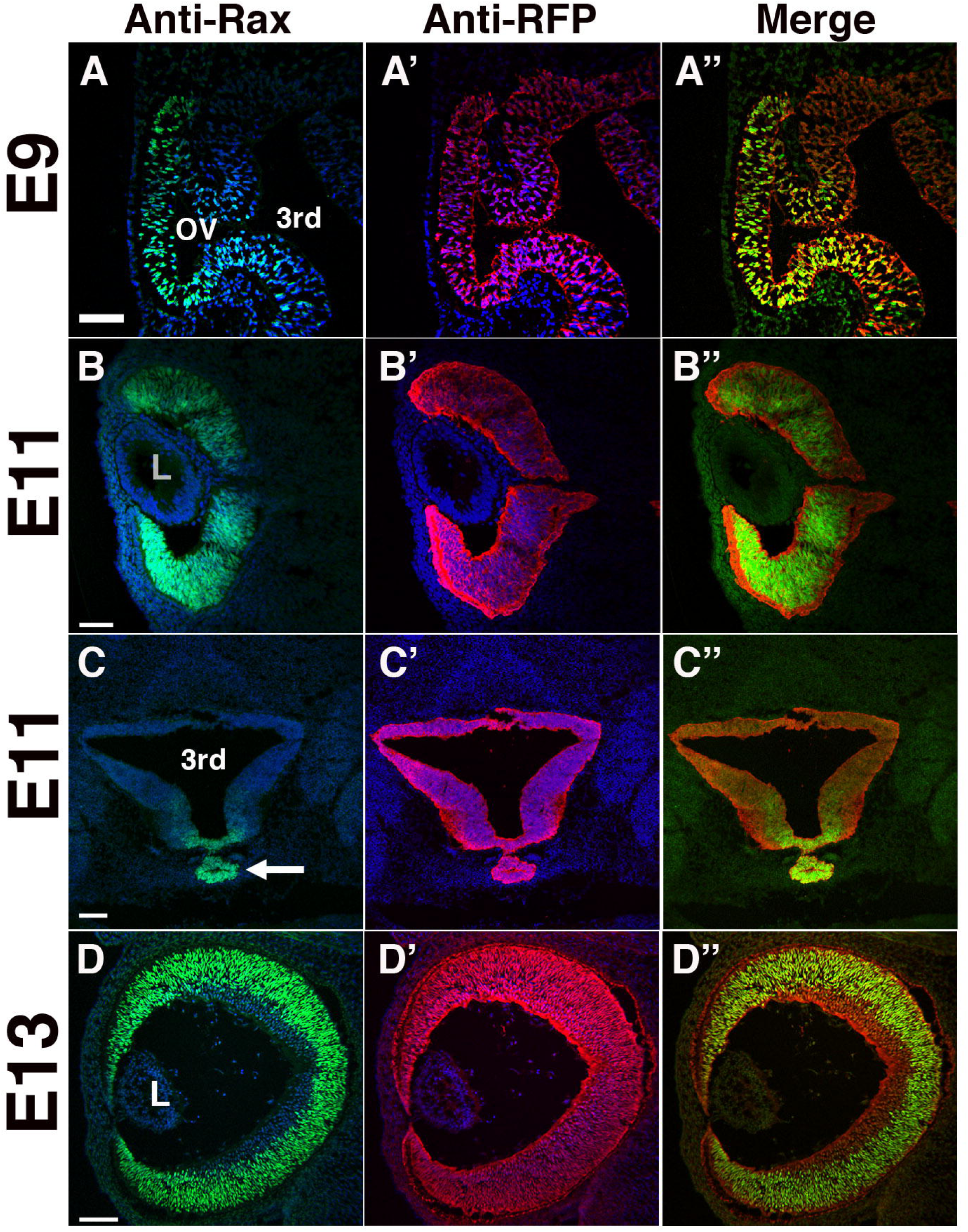
Direct comparison of endogenous Rax protein and Rax-Cre BAC Tg induced tdTomato expression. A-A”) Rax protein marks cells in the ocular field, whereas anti-DsRed labeling of tdTomato+ cells also marks adjacent brain neuroepithelial cells. **B-B”)** Rax protein is expressed by retinal progenitor cells, but Anti-DsRed also labels nascent RPE. **C-C”)** Horizontal section of diencephalon and adjacent Rathke’s pouch (arrow in C). **D-D”)** Rax nuclear protein is confined to retinal progenitor cells, whereas the tdTomato lineage includes differentiated RGCs in forming GCL. OV = optic vesicle, 3rd = third ventricle, L = lens. Scale bars A = 75 microns; B = 50 microns; C = 100 microns; D = 75 microns.

We also evaluated the Rax-Cre BAC Tg lineage expression in the adult eye (Fig. 7). Here we noted expression in the ciliary body (CB) and optic cup/retinal derived layer of the iris (Fig. 7A). At higher magnification, we found all RPE, outer nuclear (ONL), inner nuclear (INL) and ganglion cell (GCL) cells uniformly express tdTomato (Fig. 7B) or GFP (Fig. 7C). Rax-Cre; Z/EG retinal sections more easily highlighted GFP expression in Müller glia (arrows in Fig. 7C), another well-known endogenous Rax expression domain^26^.

**Figure 7.**
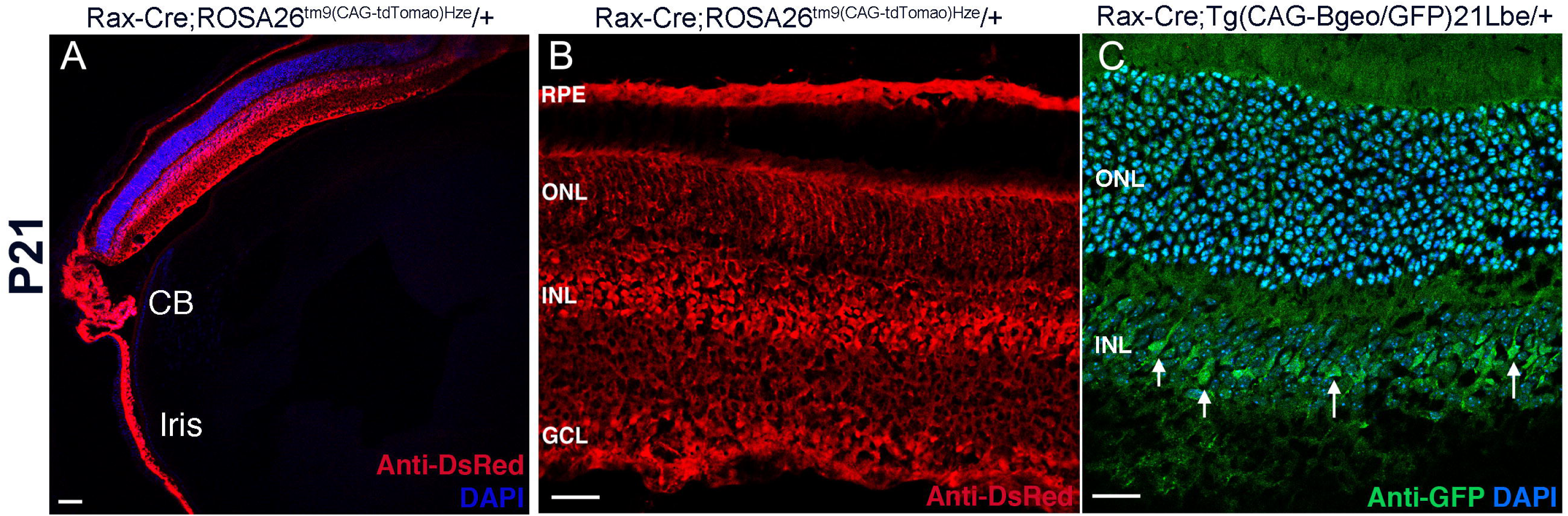
The adult Rax-Cre BAC Tg lineage. **A)** tdTomato-expressing ciliary body (CB) and iris tissues. **B)** Uniform expression of tdTomato throughout retinal layers and in RPE. **C)** Cre activated GFP is also uniformly expressed, with anti-GFP highlighting expression in Müller glia (arrows), a cell type that requires *Rax* gene activity^26^. RPE = retinal pigmented epithelium, ONL = outer nuclear layer, INL = inner nuclear layer, GCL = ganglion cell layer. Scale bars A = 50 microns; B = 50 microns; C = 100 microns.

Although Rax-Cre BAC transgenic activity is extremely faithful to endogenous Rax expression especially in the eye, we also consistently saw one ectopic limb domain (Fig. 8). From E13 to birth, all four limbs of Rax-Cre BAC Tg;Ai9 embryos have two expression subdomains (arrows in Fig. 8A, 8D,D’). Broader tdTomato expression in limb buds appears to coalesce to the first and fifth digits (Fig. 8,8D’).

**Figure 8.**
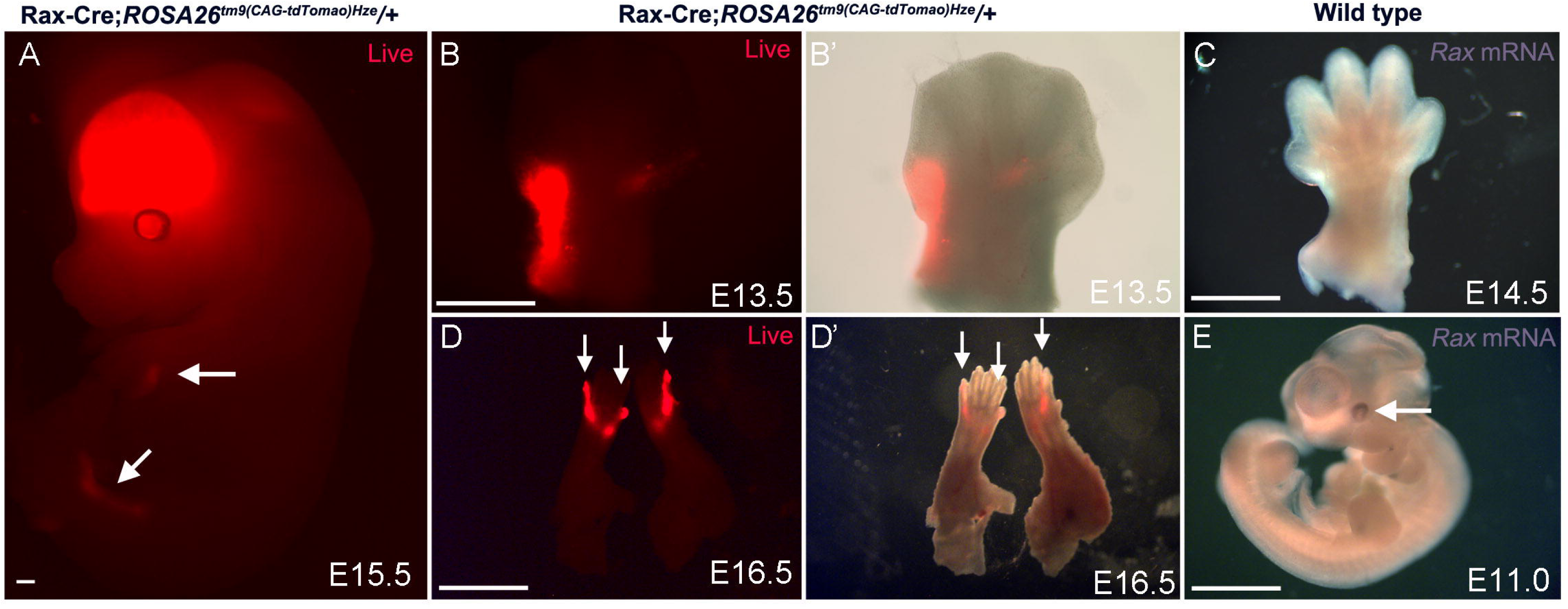
Rax-Cre BAC Tg expression during limb bud development. **A)** Cre activated tdTomato live fluorescence in the E15.5 brain, eye, forelimbs and hindlimbs (arrows). **B,B’**) tdTomato is obvious in two domains within the E13.5 left forelimb, one along the peripheral 5th digit and the other in forming 1st digit. **C**) Whole mount in site of E14.5 right forelimb showing lack of endogenous *Rax* mRNA expression. **D,D’**) Live imaging of isolated E16.5 forelimbs showing expression in the 1st and 5th forelimb digits. Hindlimbs display the same expression pattern (not shown). Panels B’ and D’ are fluorescent bright field image overlays. **E**) E11.5 whole embryo in situ with *Rax* mRNA expression in the optic cup (arrow) but no expression in limb buds. Isolated limbs in C were treated in parallel to embryos shown in E (n = 12 sets of limbs, 7 E11 embryos for 2 independent in situs). Live embryos or limbs in A, B, D represent ≥4 biologic replicates. Scale bars A = 1mm; B,D,E = 4mm; C = 5mm.

We tested for endogenous Rax mRNA expression via in situ hybridization, using whole E11.5 embryonic eye expression as positive controls (Fig. 8E, arrow) and a mixture of all limbs from twelve E14.5 wild type embryos. We found no endogenous expression during limb development, which is consistent with zero reports for this within the mouse *Rax* gene literature. Thus, we conclude there is one Rax-Cre BAC ectopic activity domain.

## Discussion

Recombination-specific (conditional) deletion of mouse genes bypasses embryonic lethality, thereby enhancing our understanding of the multiple roles for these genes. Early mouse eye development studies have benefited from several Cre-recombinase transgenes^27–29^, which are also useful for postnatal and adult mutant analyses. The homeodomain transcription factor *Rx/Rax* is one of the earliest genes expressed in the forming retina, although it is not retinal-specific. The development and use of multiple Rax-Cre drivers in numerous publications over the past twenty years demonstrate their usefulness to retinal genetics. Cre-recombinase transgenes are imperfect tools; thus, it is important to understand the strengths and weaknesses of each one. Here, our goal was to directly compare two widely used constitutive Rax-Cre mouse transgenic lines, using the very sensitive Cre-activated lineage reporter, Ai9^15^. Our datasets include live and fixed embryos across the initial stages of eye development, and cross-sections from these embryos, where Cre-activated tdTomato expression was analyzed at the cellular level. We observed novel ectopic expression domains confirming the necessity of thorough interrogation of Cre drivers.

Rx3-Cre has been widely used in dozens of functional studies for ocular, telencephalic and oral ectoderm genes (MGI database: www.informatics.jax.org). Previous studies documented ectopic Rx3-Cre activity in the surface ectoderm and differentiating lens. Our data shows additional ectopic Cre activity in the POM, starting at the optic vesicle stage. We could identify TdTomato-positive POM subpopulations lining the optic cup by their coexpression with PITX2-(neural crest-derived mesenchyme) and PECAM-1 (endothelial precursor mesodermal cells), respectively. Since these POM subpopulations are premigratory and migratory in tight association with the basal side of the RPE, the conditional loss of of critical developmental genes in these cells may influence directly cellular differentiation in choroid, sclera, RPE and anterior segment structures. For example, the transcription factor SRY (sex determining region y)-box 9 (*Sox9*) is expressed in both RPE and neural crest-derived POM^30, 31^. In the RPE *Sox9* directs a late differentiation program, but in the cranial mesenchyme, *Sox9* is required for both premigratory and migratory cranial neural crest cells^30, 32^. Thus, it is possible that Rx3-Cre;*Sox9^CKO/CKO^* mutants could exhibit both RPE and POM phenotypes, complicating a cell autonomous assignment of its requirements in either tissue. Furthermore, our own studies and others have shown a critical non-cell autonomous requirement of POM during eye development in chick and mouse^33^. We investigated the role of the O’acyltransferase *Porcn* that is required, and mostly dedicated, for posttranslational modification of Wnt ligands^18^. Using a combination of the two Cre drivers Rx3-Cre and Wnt1-Cre we showed that *Porcn* must be removed from NR, RPE, lens and POM to fully recapitulate ocular phenotypes obtained by disrupting Wnt pathway components as shown in previous studies^34, 35^. Thus, depending on the extracellular factors, dysregulated POM contribution could significantly impact differentiation and morphogenetic processes of optic cup tissues, in a non-cell autonomous manner.

For the Rax-Cre BAC transgenic, we see it onsets at E8 in the optic sulcus and expected expression in all tissues that derive from the E9 optic vesicle. We used two distinct lineage reporters to minimize bias, and they show strong, uniform expression in the forming optic cup, RPE, and optic stalk prior to the onset of retinal neurogenesis. At older developmental stages, this Rax-Cre lineage includes all retinal progenitors and the differentiating retinal cell types. In the adult retina, every retinal cell type, the RPE, ciliary body, retinal-derived iris and optic nerve each have tdTomato expression. Except for one ectopic domain (see below) we detected Cre-induced lineage reporter expression in the posterior pituitary, cortical, hypothalamic and hippocampal brain domains, as predicted. We conclude this is a validated Rax-Cre driver for constitutive conditional mutant studies of the developing visual system.

Finally, the specific expression of Rax-Cre induced tdTomato in all developing mouse limbs was unexpected (Fig. 8). The initially broad subdomains refined to just the first and fifth digits of each paw. Since *Rax* mRNA could not be detected at all in the limbs, we conclude this expression is ectopic.

Interestingly, *Rax/RAX* genes have been duplicated or lost several times during vertebrate evolution^36^. Humans genomes contain *RAX 1* and *RAX 2* genes that are separately required for normal vision^37^, whereas mice have a single *Rax* gene. Evolutionarily, invertebrate genomes contain a gene cluster with three PRD-class homeobox genes: *homeobrain*, *rx* and *orthopedia* (Otp). The expression and regulation of these genes have been studied in multiple cnidarians, insect and mollusk species^38^. Relevant here is that there is coordinate expression in the appendages (e.g., *Nemastella* tentacles)^38^. Yet in vertebrates, this gene cluster is no longer identifiable and the *homeobrain* gene is missing. *Rx/Rax* and *Otp* genes frequently reside on different chromosomes, with only Rax present in the eye, while Rax and Otp are each required in the particular tissues of the developing brain^25, 39^. Thus, it theoretically possible that *Rax* or *Otp* genes retain an ancient, but silenced, appendage enhancer; with Rax transgenics, by virtue of their independent genomic insertions, removing local silencing constraints on this enhancer. More extensive, comparative genomics and cross-species experiments are needed to better support this idea. Overall, it is important to gain deeper understanding of *RAX* gene regulation, particularly if its regulatory regions would be used to target therapeutics for treating ocular disease.

In summary, in this study we provide a systematic analysis of two widely utilized Rax-Cre drivers during embryonic and postnatal eye development. Importantly, these Cre drivers are very useful to recapitulate severe congenital eye diseases as shown for Rx3-Cre, for example Charge syndrome, coloboma, Focal Dermal Hypoplasia and Leber congenital amaurosis (MGI database: www.informatics.jax.org).

## Acknowledgements

The authors thank Nadege Lum Ellerman, Kelly McCulloh, Sarah Kiser, Emelie McKenzie and Yang Song for technical support; Tom Glaser, Anna La Torre and Edward Levine for critical feedback and discussion.

---

**Suppl. Figure 1. Pedigree showing transmission of germ line deletion of *Taz*.** Rx3-Cre originated from a single female breeder #4 in generation I on a mixed wild type background and was inherited by son #2 in generation II (mixed wild type). Progeny in generation III inherited both Rx3-Cre and showed partially excision of the single conditional allele. In generation IV, occurrence of wild type males and females with or with Rx3-Cre are shown with no indication of partially or fully excised *Taz* (filled black symbols, #1,4), a male with Cre and full homozygous excision (open square, #6), three progeny with Rx3-Cre and partially or not excised Taz alleles (#2,7,10). The remaining progeny harbor one fully excised *Taz* allele with Cre (#3,5) and, interestingly, without Rx3-Cre (#8,9). Symbols: circles are females, squares are male; see key for more explanation.

## References

1. Diacou R, Nandigrami P, Fiser A, Liu W, Ashery-Padan R, Cvekl A. Cell fate decisions, transcription factors and signaling during early retinal development. Progress in retinal and eye research 2022;91:101093.

2. Miesfeld JB, Brown NL. Eye organogenesis: A hierarchical view of ocular development. Current topics in developmental biology 2019;132:351–393.

3. Furukawa T, Kozak CA, Cepko CL. rax, a novel paired-type homeobox gene, shows expression in the anterior neural fold and developing retina. Proceedings of the National Academy of Sciences of the United States of America 1997;94:3088–3093.

4. Mathers PH, Grinberg A, Mahon KA, Jamrich M. The Rx homeobox gene is essential for vertebrate eye development. Nature 1997;387:603–607.

5. Brachet C, Kozhemyakina EA, Boros E, et al. Truncating RAX Mutations: Anophthalmia, Hypopituitarism, Diabetes Insipidus, and Cleft Palate in Mice and Men. J Clin Endocrinol Metab 2019;104:2925–2930.

6. Davis ES, Voss G, Miesfeld JB, Zarate-Sanchez J, Voss SR, Glaser T. The rax homeobox gene is mutated in the eyeless axolotl, Ambystoma mexicanum. Developmental dynamics: an official publication of the American Association of Anatomists 2021;250:807–821.

7. Voronina VA, Kozhemyakina EA, O’Kernick CM, et al. Mutations in the human RAX homeobox gene in a patient with anophthalmia and sclerocornea. Human molecular genetics 2004;13:315–322.

8. Zhang L, Mathers PH, Jamrich M. Function of Rx, but not Pax6, is essential for the formation of retinal progenitor cells in mice. Genesis 2000;28:135–142.

9. Stigloher C, Chapouton P, Adolf B, Bally-Cuif L. Identification of neural progenitor pools by E(Spl) factors in the embryonic and adult brain. Brain Res Bull 2008;75:266– 273.

10. Swindell EC, Bailey TJ, Loosli F, et al. Rx-Cre, a tool for inactivation of gene expression in the developing retina. Genesis 2006;44:361–363.

11. Heintz N. Gene expression nervous system atlas (GENSAT). Nature neuroscience 2004;7:483.

12. Klimova L, Lachova J, Machon O, Sedlacek R, Kozmik Z. Generation of mRx-Cre Transgenic Mouse Line for Efficient Conditional Gene Deletion in Early Retinal Progenitors. PloS one 2013;8:e63029.

13. Bosze B, Moon MS, Kageyama R, Brown NL. Simultaneous Requirements for Hes1 in Retinal Neurogenesis and Optic Cup-Stalk Boundary Maintenance. The Journal of neuroscience: the official journal of the Society for Neuroscience 2020;40:1501– 1513.

14. Bosze B, Suarez-Navarro J, Cajias I, Brzezinski Iv JA, Brown NL. Notch pathway mutants do not equivalently perturb mouse embryonic retinal development. PLoS genetics 2023;19:e1010928.

15. Madisen L, Garner AR, Shimaoka D, et al. Transgenic mice for intersectional targeting of neural sensors and effectors with high specificity and performance. Neuron 2015;85:942–958.

16. Brown NL, Kanekar S, Vetter ML, Tucker PK, Gemza DL, Glaser T. Math5 encodes a murine basic helix-loop-helix transcription factor expressed during early stages of retinal neurogenesis. Development 1998;125:4821–4833.

17. Tucker RP, Chiquet-Ehrismann R, Chevron MP, Martin D, Hall RJ, Rubin BP. Teneurin-2 is expressed in tissues that regulate limb and somite pattern formation and is induced in vitro and in situ by FGF8. Developmental dynamics: an official publication of the American Association of Anatomists 2001;220:27–39.

18. Bankhead EJ, Colasanto MP, Dyorich KM, Jamrich M, Murtaugh LC, Fuhrmann S. Multiple requirements of the focal dermal hypoplasia gene porcupine during ocular morphogenesis. Am J Pathol 2015;185:197–213.

19. Sun WR, Ramirez S, Spiller KE, Zhao Y, Fuhrmann S. Nf2 fine-tunes proliferation and tissue alignment during closure of the optic fissure in the embryonic mouse eye. Human molecular genetics 2020;29:3373–3387.

20. Gage PJ, Rhoades W, Prucka SK, Hjalt T. Fate maps of neural crest and mesoderm in the mammalian eye. Investigative ophthalmology & visual science 2005;46:4200–4208.

21. Lwigale PY. Corneal Development: Different Cells from a Common Progenitor. Prog Mol Biol Transl Sci 2015;134:43–59.

22. Walls JR, Coultas L, Rossant J, Henkelman RM. Three-dimensional analysis of vascular development in the mouse embryo. PloS one 2008;3:e2853.

23. Lu F, Kar D, Gruenig N, et al. Rax is a selector gene for mediobasal hypothalamic cell types. The Journal of neuroscience: the official journal of the Society for Neuroscience 2013;33:259–272.

24. Muranishi Y, Terada K, Furukawa T. An essential role for Rax in retina and neuroendocrine system development. Dev Growth Differ 2012;54:341–348.

25. Orquera DP, Nasif S, Low MJ, Rubinstein M, de Souza FSJ. Essential function of the transcription factor Rax in the early patterning of the mammalian hypothalamus. Developmental biology 2016;416:212–224.

26. Furukawa T, Mukherjee S, Bao ZZ, Morrow EM, Cepko CL. rax, Hes1, and notch1 promote the formation of Muller glia by postnatal retinal progenitor cells. Neuron 2000;26:383–394.

27. Furuta Y, Lagutin O, Hogan BL, Oliver GC. Retina- and ventral forebrain-specific Cre recombinase activity in transgenic mice. Genesis 2000;26:130–132.

28. Marquardt T, Ashery-Padan R, Andrejewski N, Scardigli R, Guillemot F, Gruss P. Pax6 is required for the multipotent state of retinal progenitor cells. Cell 2001;105:43– 55.

29. Muranishi Y, Terada K, Inoue T, et al. An essential role for RAX homeoprotein and NOTCH-HES signaling in Otx2 expression in embryonic retinal photoreceptor cell fate determination. The Journal of neuroscience: the official journal of the Society for Neuroscience 2011;31:16792–16807.

30. Cohen-Tayar Y, Cohen H, Mitiagin Y, et al. Pax6 regulation of Sox9 in the mouse retinal pigmented epithelium controls its timely differentiation and choroid vasculature development. Development 2018;145.

31. Kim JY, Park R, Lee JH, et al. Yap is essential for retinal progenitor cell cycle progression and RPE cell fate acquisition in the developing mouse eye. Developmental biology 2016;419:336–347.

32. Barrionuevo F, Bagheri-Fam S, Klattig J, et al. Homozygous inactivation of Sox9 causes complete XY sex reversal in mice. Biol Reprod 2006;74:195–201.

33. Fuhrmann S. Eye morphogenesis and patterning of the optic vesicle. Current topics in developmental biology 2010;93:61–84.

34. Alldredge A, Fuhrmann S. Loss of Axin2 Causes Ocular Defects During Mouse Eye Development. Investigative ophthalmology & visual science 2016;57:5253–5262.

35. Fuhrmann S. Wnt signaling in eye organogenesis. Organogenesis 2008;4:60–67.

36. Kon T, Furukawa T. Origin and evolution of the Rax homeobox gene by comprehensive evolutionary analysis. FEBS Open Bio 2020;10:657–673.

37. Orquera DP, de Souza FSJ. Evolution of the Rax family of developmental transcription factors in vertebrates. Mechanisms of development 2017;144:163–170.

38. Mazza ME, Pang K, Reitzel AM, Martindale MQ, Finnerty JR. A conserved cluster of three PRD-class homeobox genes (homeobrain, rx and orthopedia) in the Cnidaria and Protostomia. Evodevo 2010;1:3.

39. Simeone A, D’Apice MR, Nigro V, et al. Orthopedia, a novel homeobox-containing gene expressed in the developing CNS of both mouse and Drosophila. Neuron 1994;13:83–101.

