## Supplementary figures and images for "Direct comparison of constitutive Rax-Cre transgenic drivers that activate in the mouse embryonic eye field"

### Supplemental Figure 1

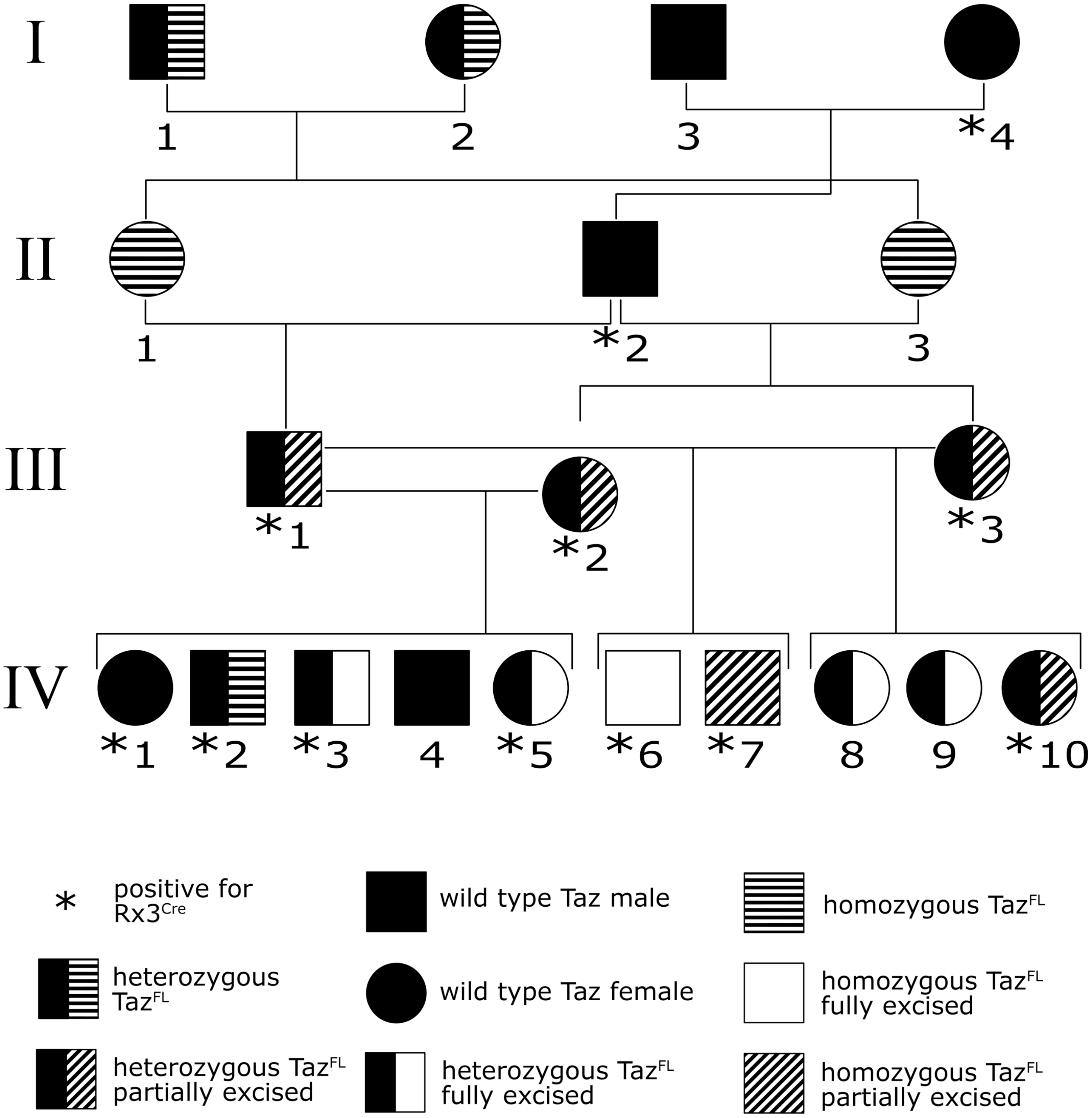
